# Using the Weibull accelerated failure time regression model to predict time to health events

**DOI:** 10.1101/362186

**Authors:** Enwu Liu, Karen Lim

## Abstract

We describe a statistical method protocol to use a Weibull accelerated failure time (AFT) model to predict time to a health-related event. This prediction method is quite common in engineering reliability research but rarely used for medical predictions such as survival time. A worked example for how to perform the prediction using a published dataset is included.

## Introduction

In medical literature, most prediction models are used to predict the probability of an event occurring or condition developing over a specified time period, such as the Framingham 10 Year Risk of General Cardiovascular Disease [1] and FRAX: a Tool for Estimating 10 year Fracture Risk. [2] In engineering reliability research, it is common to use Weibull accelerated failure time (AFT) models to predict ‘time to failure’, for example the lifespan of a machine; when a component will need replacement; and an optimal maintenance schedule that maximizes reliability of the entire system. [3] Weibull AFT models are also used to predict shelf-life of perishable foods and warranty period of goods. [4, 5] This statistical method estimates when the event will occur without being bound to a defined time period (i.e. absolute time; when a component will need replacement vs. relative time; 10-y risk of needing replacement). In addition to predicting engineering or mechanic events, this statistical method might also be useful for predicting medical events such as fracture, myocardial infarction and death.

In this paper, we have not intended to develop and present a prediction tool. Rather, we aim to illustrate how to use the Weibull AFT model and assess point accuracy in a medical context.

### Weibull distribution

The Weibull distribution is also called the type III extreme value distribution. [6] The distribution has three parameters: the location parameter *μ*, the scale parameter *ρ* and the shape parameter *γ*. The location parameter *μ* is predetermined as the minimum value in the distribution. In survival or failure analysis, *μ* of 0 usually selected to produce a two-parameter distribution.

The cumulative distribution function (CDF) for a two-parameter Weibull distributed random variable *T ∼ W* (*ρ, γ*) is given as

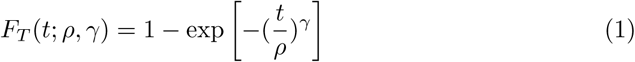

for *t ≥* 0, *ρ >* 0, *γ >* 0 and *F* (*t*; *ρ, γ*) = 0 for *t <* 0

The probability density function (PDF) of the Weibull distribution is given as

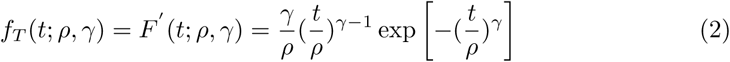

The survival function of the Weibull distribution is given as

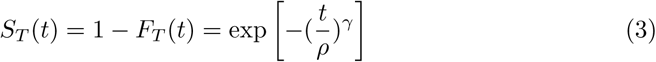

The mean survival time or mean time to failure (MTTF) is given as

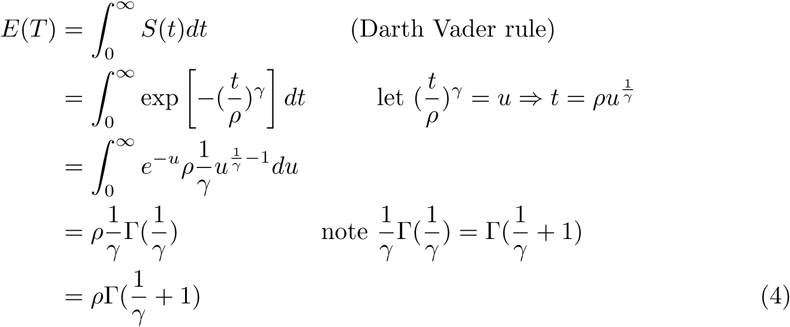

### Log-Weibull distribution

The log-Weibull distribution is also called Gumbel distribution, or type I extreme value distribution. [7] Let *T ∼ W* (*ρ, γ*) and *Y* = *g*(*T*) = *logT* is a one-one transformation from support *𝒯* = {*t|t >* 0} to *𝒴* = {*y| - ∞ < y < ∞*}. The inverse of *Y* is

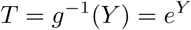

and the Jacobian is given as

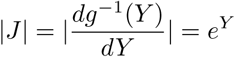

The probability density function of *Y* is then derived from (2)

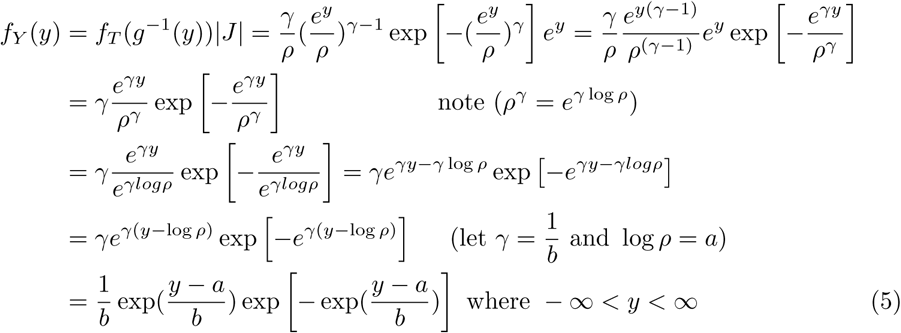

It shows log-Weibull distribution has a Gumbel distribution *G*(*a, b*), where *a* = log *ρ* and 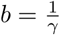. The CDF of the log-Weibull distribution can be derived from the above PDF or by the definition directly.

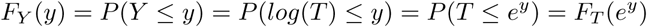

By Eq(1) we get

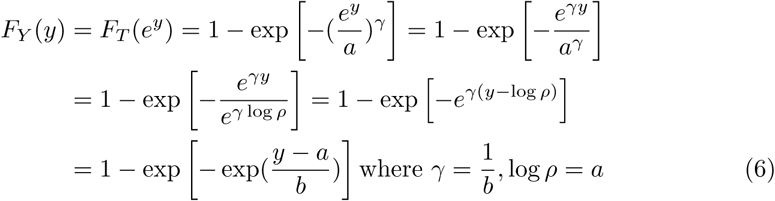

The survival function of *Y* = log(*T*) is given by

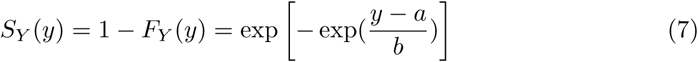

The hazard function is given by

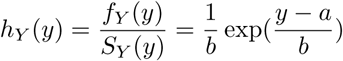

### Weibull AFT regression model

Let *T* be the survival time. Suppose we have a random sample of size *n* from a target population. For a subject *i*(*i* = 1, 2, *…, n*), we have observed values of covariates *x*_*i*1_, *x*_*i*2_, *…, x*_*ip*_ and possibly censored survival time *t*_*i*_. We can express the Weibull AFT model as

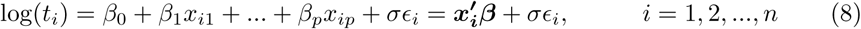

where ***β*** = (*β*_0_, *…, β*_*p*_) are the regression coefficients of interest, *σ* is a scale parameter and *ϵ*_1_, *… ϵ*_*n*_ are independent and identically distributed according to a Gumbel distribution with PDF

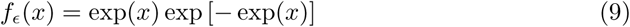

and the CDF

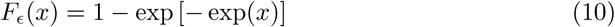

Note Eq(9) and Eq(10) are equal to set *a* = 0, *b* = 1 in Eq(5) and Eq(6). We denote it as a *G*(0, 1) distribution or a standard Gumbel distribution.

Now, let us find the PDF of *T* from Eq(8)

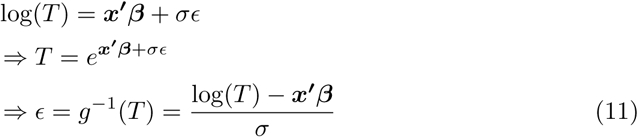

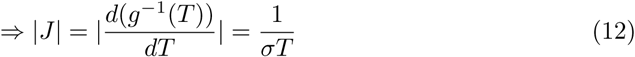

Puting Eq(11) and Eq(12) to Eq(9) we get

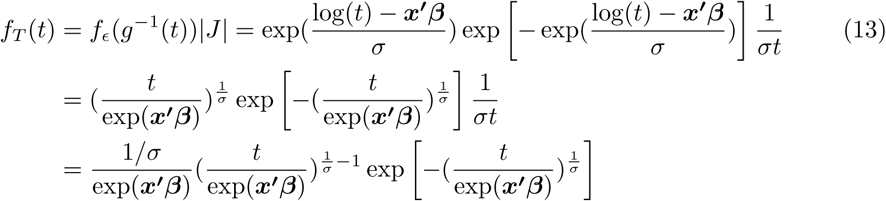

Comparing Eq(13) with Eq (2) and let 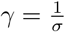 and *ρ* = exp(***x*^′^*β***), we can see *T* has a Weibull distribution 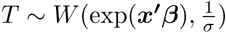.

Referring to Eq(3), now the survival function of 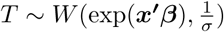 can be written as

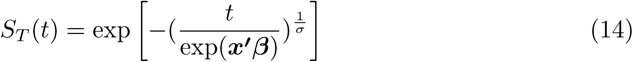

Referring to Eq(4) the expected survival time of 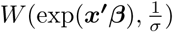 is given as

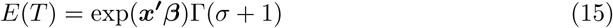

Since most statistical software use log(*T*) to calculate these parameters, let us show the distribution and characteristics of the log(*T*). Let

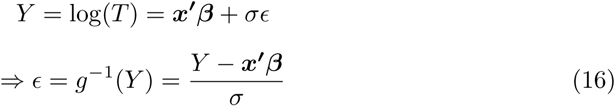

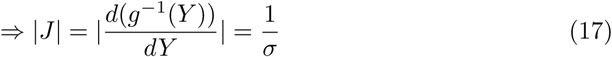

Putting Eq(16) and Eq(17) to Eq(9), we get

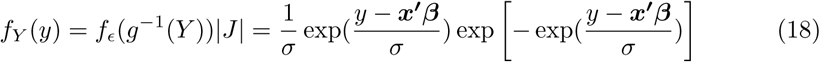

Comparing Eq(18) to Eq(5) we can see that *Y* i.e. log(*T*) has a *G*(***x*^′^*β***, *σ*) distribution. We also can see the error term *ϵ*, which has a *G*(0, 1) distribution. similar to the error term in a simple linear regression with a *N* (0, *σ*^2^) distribution.

Referring to Eq(13) and Eq(18), we can see in the Weibull AFT model, *T* has a Weibull 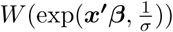 distribution, and log(*T*) has a Gumbel *G*(***x*^′^*β***, *σ*) distribution.

From Eq (7) the survival function of *Y* i.e. log(*T*) is given as

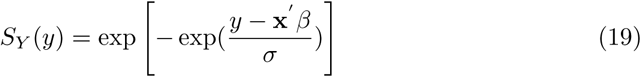

And the expectation of *Y* i.e log(*T*) is calculated as

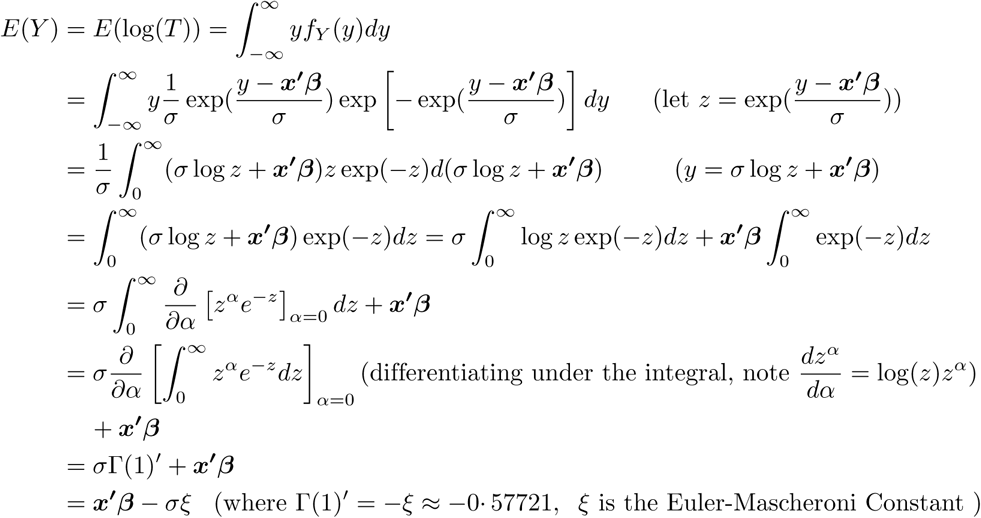

Note, since log(*x*) is a concave down function by Jensen’s inequality, *E*(log(*T*)) *≤* log(*E*(*T*)), Eq (15) is the correct formula to calculate the expected survival time rather than exp(***x*^′^*β*** *- σξ*).

### Parameter estimation for the Weibull AFT model

The parameters of Weibull AFT model can be estimated by the maximum likelihood method. The likelihood function of the *n* observed log(*t*) time, *y*_1_, *y*_2_, *…, y*_*n*_ is given by

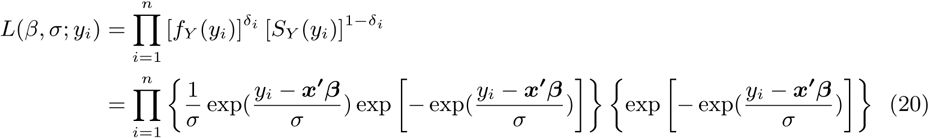

where *δ*_*i*_ is the event indicator for ith subject with *δ*_*i*_ = 1 if an event has occurred and *δ*_*i*_ = 0 if the event has not occurred. The maximum likelihood estimates *p* + 1 parameters *σ, β*_1_, *β*_*p*_. We can take the log of the likelihood function and use the Newton-Raphson method to calculate these parameters. Most statistical software packages can perform these calculations.

### Calculate expected survival time using the Weibull AFT model

In reliability research, expected survival time is called mean time to failure (MTTF) or mean time between failures (MTBF). [8]

Suppose we want to predict a person *i*’s mean survival time *t*_*i*_ using the Weibull AFT model. First, we use the maximum likelihood estimation (MLE) method of Eq(20) to calculate the 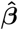 and 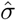 then by the invariance property of MLE, we use Eq(15) to calculate the predicted MTTF directly.

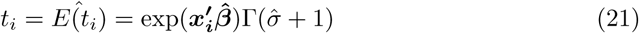

After we calculate the MTTF, we can use the Delta method to calculate the confidence interval for the MTTF. We will treat the predicted MTTF as a function 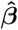 and 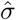. The standard error of the MTTF can be calculated as

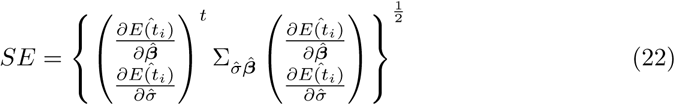

where 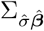 is the variance-covariance matrix of 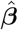 and 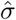. The variance-covariance matrix can be estimated by the observed Fisher information of the Weibull AFT model. The (1-*α*)% confidence interval is given as

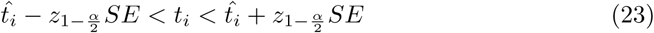

where *α* is the type I error and *z* is the quantile of the standard normal distribution.

### Calculate median survival time using Weibull AFT model

Another important statistic in survival analysis is the median survival time or percentile survival time. The *p*th percentile of survival time is calculated from the survival function. For the individual *i* the *p*th percentile of survival time is calculated as

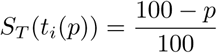

For the Weibull AFT model, we use Eq(14) to calculate *p*th survival time of an individual *i*.

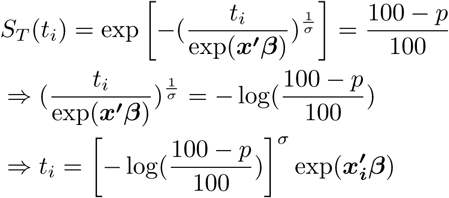

After we obtain 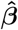 and 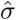 from Eq(20) and use the the invariance property of MLE, the median survival time is estimated by

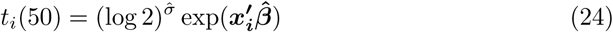

We can treat the estimated survival time percentile as a function of 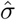 and 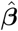 when *p* is fixed by using the Delta method to calculate the standard error of predicted *p*th survival time. The method is the same as in Eq(22) and Eq(23)

### Minimum prediction error survival time (MPET)

Both mean and median survival time estimates are biased when the sample size is small and the model includes censoring. To attenuate bias, Henderson et al. purposed a method to find the optimum prediction time with the minimum prediction error. [9] They specified that if an observed survival time *t* falls in the interval 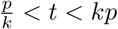 where *p* is the predicted survival time and *k >* 1 then the prediction is accurate. The probability of prediction error *E*_*k*_ condition on the predicted time *p* is given as

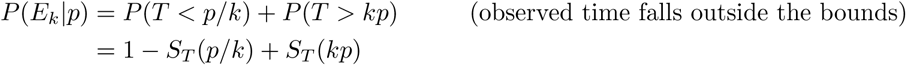

The minimum probability of prediction error *P* (*E*_*k*_*|p*) is achieved when

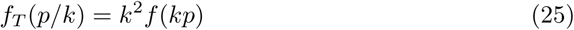

Now let us calculate the minimum prediction error for the Weibull AFT model. From Eq(13) we get

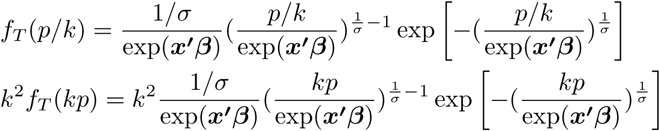

Applying Eq(25) and canceling the common parts, we get

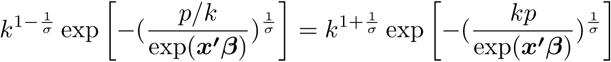

Taking the log of both sides we get

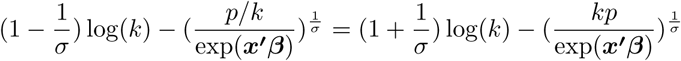

Finally, when rearranging these terms we get

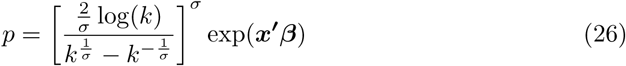

Here *p* is the minimum prediction error survival time. We may use the Delta method to get the standard error of the minimum prediction error survival time. Bootstrap methods also could be used to get a confidence interval.

### An example to predict the survival time

We use a published larynx cancer dataset of 90 male participants [10] to demonstrate the calculations of the predictions. The dataset includes five variables: stage of cancer (’stage’; 1=stage 1, 2=stage 2, 3=stage 3, 4=stage 4); time to death or on-study time in months (’time’); age at diagnosis of larynx cancer (’age’); year of diagnosis (’diagyr’); and death indicator (’delta’; 0=alive, 1=dead). The authors added a unique particiapnt identifier variable (ID) to the dataset and renamed the variable ‘delta’ to ‘death’. The dataset can be downloaded from *here*. The resulting dataset has the following structure

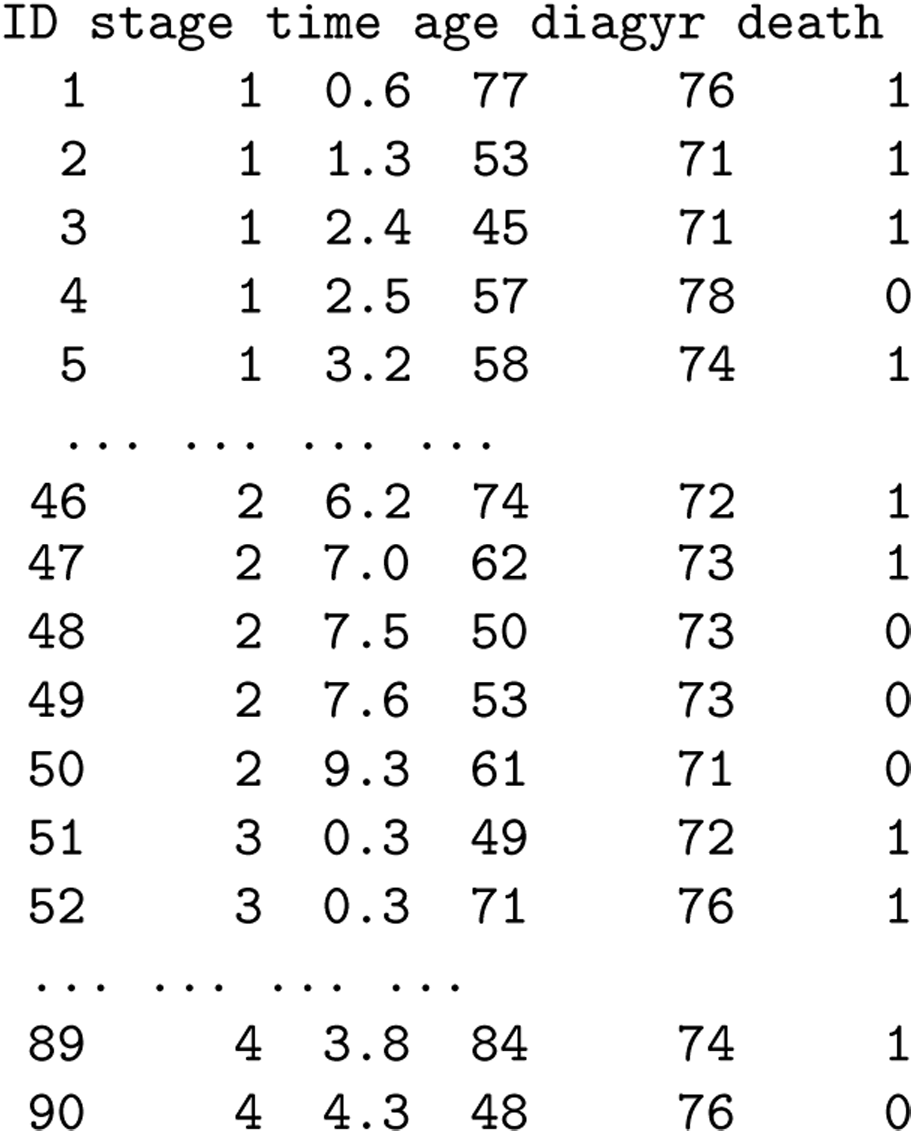

We will use two predictors to predict the survival time: 1) stage of cancer and 2) age at cancer diagnosis. Since stage of cancer is a categorical variable, we will create three dummy variables for ‘stage’ and use stage 1 as the default reference group. The Weibull AFT model can be written as

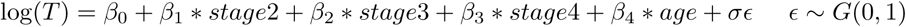

Most statistical software can run the Weibull AFT regression model. Here we display R syntax

~~~
library(survival)
larynx<-read.csv(“D:/larynx.csv”)
wr <- survreg(Surv(time, death) ∼ factor(stage) + age,
data = larynx, dist="w”)
summary(wr)
~~~

The above syntax yields the following results

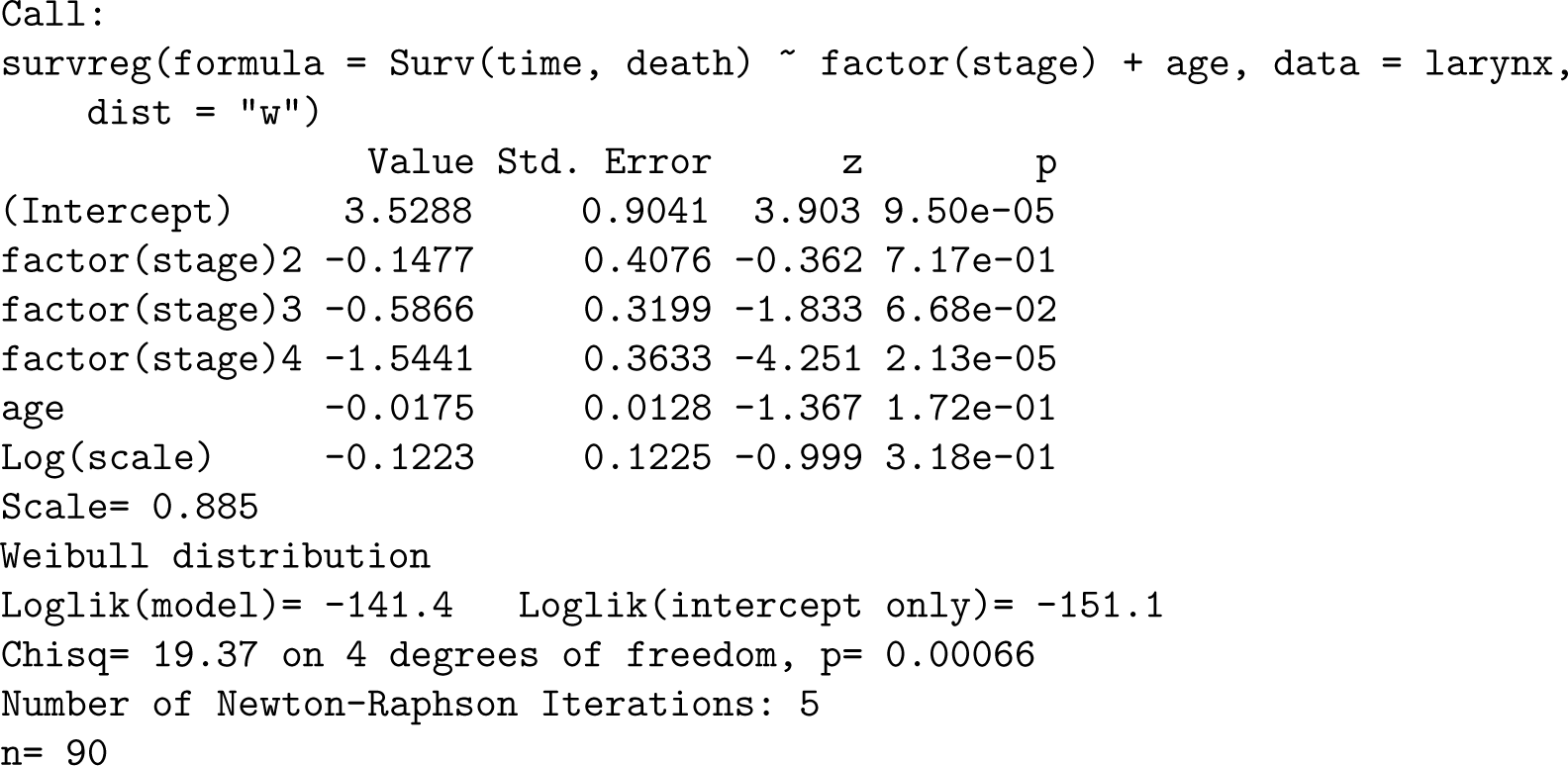

Suppose we wanted to apply all three methods (MTTF, median and MPET) to predict the survival time of patient ID=46, who had stage 2 cancer and was aged 74 years.

We would use Eq(21) to calculate his expected survival time (mean or MTTF)

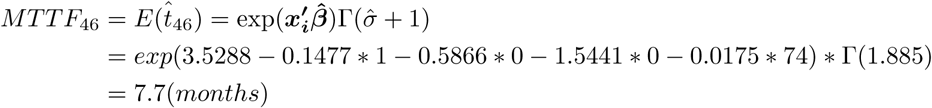

We would use Eq(24) to predict his median survival time

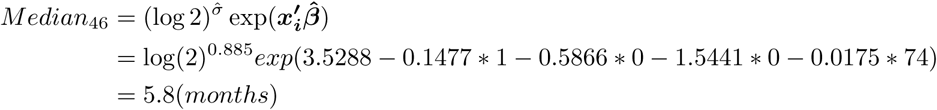

We would use Eq(26) to calculate the MPET and we fix *k*=2.

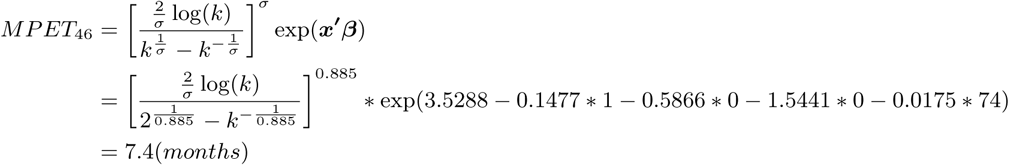

It seems the three prediction methods MTTF, median and MPET produce estimates close to the actual survival time of patient ID=46, which was 6.2 months.

Note in R build in *predict* function for Weibull AFT model type= “response” calculates exp 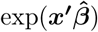 without the 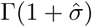 and type=”lp” calculates 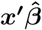 only. We should not use them to predict MTTF and there are no software to calculate the minimum prediction error survival time

We use the Delta method to calculate the 95% confidence interval of the median survival time. The standard error of the predicted median time is calculated using Eq(22)

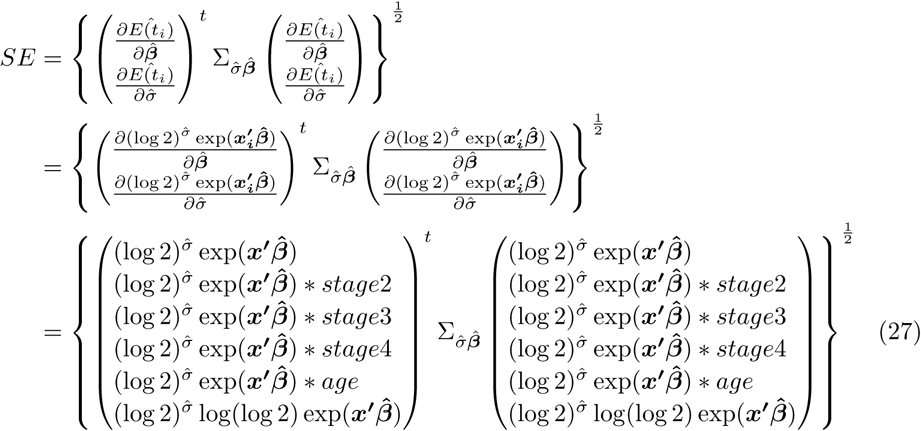

First, we need to find the variance-covariance matrix 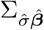, which can be calculated by the observed Fisher information of the Weibull AFT model (calculated by most software). Again in R:

`wr$var`

We get

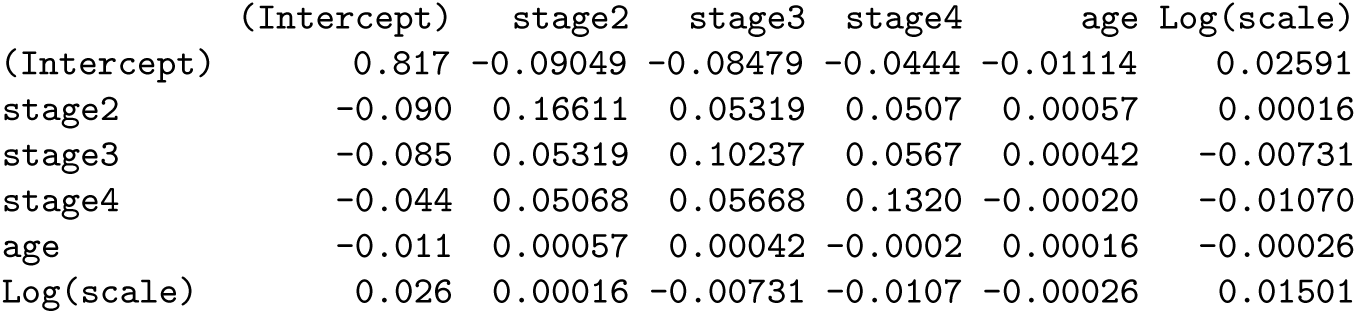

Note, R generates the variance-covariance of coefficients as log(scale), so we need to change the log(scale) to scale by performing extra calculations. Using formulas on page 401 of the book by John Klein [11], set *g*_1_(*σ,* ***β***) = ***β*** and *g*_2_(*σ,* ***β***) = *σ* = *e*^log(*σ*)^, i.e *σ* = *e*^*θ*^ is a function of *θ* and *θ* = log(*σ*), we get

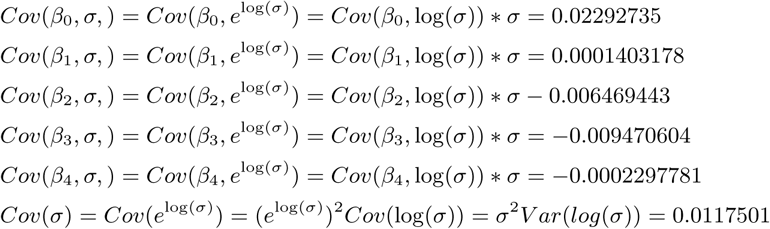

Now we will use these six values to replace the last row and column of the variance-covariance matrix of the coefficients and log(scale). We get

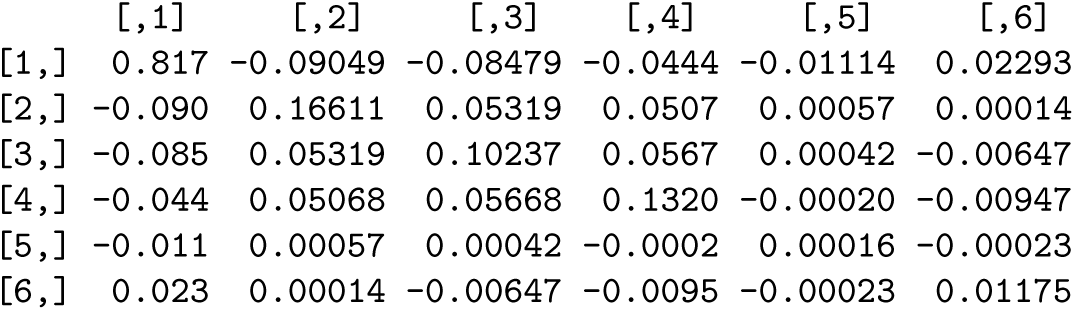

This is the 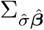 matrix we needed.

If we use SAS software, we can directly get the variance-covariance matrix of 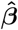 and 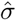 by using the following statements

~~~
proc lifereg data=larynx order=data COVOUT outest=est;
class stage;
model time*death(0)=stage age/dist=weibull;
run;
proc print data=est;
run;
~~~

The right side vector of the Eq(27) is calculated as

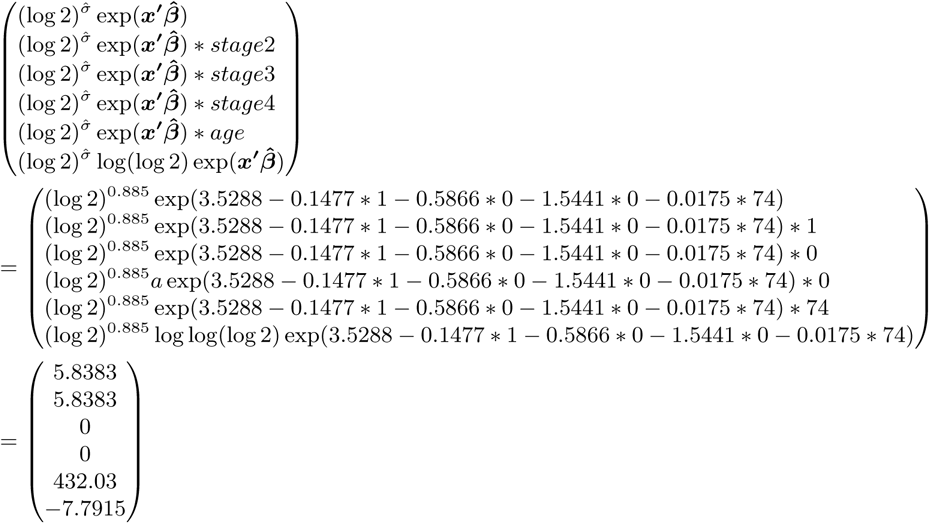

Now we have everything to calculate the standard error of the median survival time.

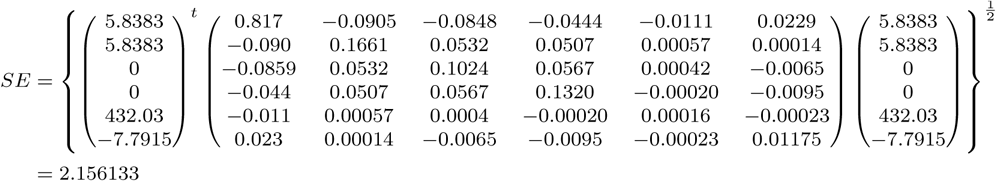

Therefore, the 95% confidence interval is given as

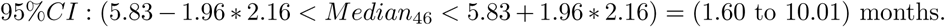

This result indicates we are 95% confident that the survival time will be within 1.60 to 10.01 months. We can also use the R build in function *predict* to predict the median survival time.

~~~
Median46<-predict(wr, newdata=data.frame(stage=2, age=74), type="quantile", p=0.5, se.fit=TRUE)
Median46
~~~

We get

~~~
$fit
  5.838288
$se.fit
2.095133
~~~

This standard error differs slightly to our calculations as R uses Greedwood’s formula to calculate the standard error of the survival function. [11]

### Assess the point prediction accuracy

Two methods of assessing accuracy of predicted survival time have been proposed by Parkes and Christakis and Lamont. [12, 13] Parkes suggests let *t* be the observed survival time and *p* be the predicted time. If *p/k ≤ t ≤ kp* then the point prediction *p* is defined as “accurate” and any value outside the interval is “inaccurate”. Parkes proposes *k* = 2 as suitable. As a more strict assessment, Christakis and Lamont purposed a 33% rule to measure accuracy, where the observed time is divided by the predicted survival time and defined as “accurate” if this quotient lies between 0.67 and 1.33. Values less than or greater than 1.33 are defined as “errors”. We compared the accuracy of our example using Parkes method (*k* = 2) and Christakis’ method. The accuracy rate (i.e. proportion of “accurate” prediction over the total sample size) is presented in Table 1.

**Table 1.** Prediction results and accuracy (last digit in predicted time: 0 inaccurate, 1, accurate)

| ID | stage | age | death | time | Median(95% CI) | MTTF | MPET |
| --- | --- | --- | --- | --- | --- | --- | --- |
| 1 | 1 | 77 | 1 | 0.6 | 6.42(3.16,9.68),0 | 8.47,0 | 8.11,0 |
| 2 | 1 | 53 | 1 | 1.3 | 9.77(4.07,15.46),0 | 12.9,0 | 12.34,0 |
| 3 | 1 | 45 | 1 | 2.4 | 11.23(3.07,19.39),0 | 14.84,0 | 14.19,0 |
| 4 | 1 | 57 | 0 | 2.5 | 9.11(4.32,13.89),0 | 12.03,0 | 11.5,0 |
| 5 | 1 | 58 | 1 | 3.2 | 8.95(4.36,13.54),0 | 11.82,0 | 11.3,0 |
| 6 | 1 | 51 | 0 | 3.2 | 10.11(3.88,16.34),0 | 13.36,0 | 12.78,0 |
| 7 | 1 | 76 | 1 | 3.3 | 6.54(3.29,9.78),1 | 8.62,0 | 8.25,0 |
| 8 | 1 | 63 | 0 | 3.3 | 8.2(4.37,12.03),0 | 10.83,0 | 10.36,0 |
| 9 | 1 | 43 | 1 | 3.5 | 11.63(2.72,20.54),0 | 15.36,0 | 14.7,0 |
| 10 | 1 | 60 | 1 | 3.5 | 8.64(4.4,12.89),0 | 11.41,0 | 10.91,0 |
| 11 | 1 | 52 | 1 | 4 | 9.94(3.98,15.89),0 | 13.13,0 | 12.55,0 |
| 12 | 1 | 63 | 1 | 4 | 8.2(4.37,12.03),0 | 10.83,0 | 10.36,0 |
| 13 | 1 | 86 | 1 | 4.3 | 5.49(1.96,9.01),1 | 7.24,1 | 6.92,1 |
| 14 | 1 | 48 | 0 | 4.5 | 10.66(3.52,17.79),0 | 14.08,0 | 13.47,0 |
| 15 | 1 | 68 | 0 | 4.5 | 7.52(4.12,10.91),1 | 9.92,0 | 9.49,0 |
| 16 | 1 | 81 | 1 | 5.3 | 5.99(2.63,9.34),1 | 7.9,1 | 7.56,1 |
| 17 | 1 | 70 | 0 | 5.5 | 7.26(3.96,10.56),1 | 9.58,1 | 9.16,1 |
| 18 | 1 | 58 | 0 | 5.9 | 8.95(4.36,13.54),1 | 11.82,0 | 11.3,1 |
| 19 | 1 | 47 | 0 | 5.9 | 10.84(3.38,18.31),1 | 14.33,0 | 13.7,0 |
| 20 | 1 | 75 | 1 | 6 | 6.65(3.41,9.89),1 | 8.78,1 | 8.39,1 |
| 21 | 1 | 77 | 0 | 6.1 | 6.42(3.16,9.68),1 | 8.47,1 | 8.11,1 |
| 22 | 1 | 64 | 0 | 6.2 | 8.06(4.34,11.77),1 | 10.64,1 | 10.18,1 |
| 23 | 1 | 77 | 1 | 6.4 | 6.42(3.16,9.68),1 | 8.47,1 | 8.11,1 |
| 24 | 1 | 67 | 1 | 6.5 | 7.65(4.19,11.1),1 | 10.1,1 | 9.66,1 |
| 25 | 1 | 79 | 0 | 6.5 | 6.2(2.9,9.5),1 | 8.18,1 | 7.83,1 |
| 26 | 1 | 61 | 0 | 6.7 | 8.49(4.4,12.58),1 | 11.21,1 | 10.73,1 |
| 27 | 1 | 66 | 0 | 7 | 7.78(4.25,11.31),1 | 10.27,1 | 9.83,1 |
| 28 | 1 | 68 | 1 | 7.4 | 7.52(4.12,10.91),1 | 9.92,1 | 9.49,1 |
| 29 | 1 | 73 | 0 | 7.4 | 6.89(3.65,10.12),1 | 9.09,1 | 8.69,1 |
| 30 | 1 | 56 | 0 | 8.1 | 9.27(4.28,14.26),1 | 12.24,1 | 11.71,1 |
| 31 | 1 | 73 | 0 | 8.1 | 6.89(3.65,10.12),1 | 9.09,1 | 8.69,1 |
| 32 | 1 | 58 | 0 | 9.6 | 8.95(4.36,13.54),1 | 11.82,1 | 11.3,1 |
| 33 | 1 | 68 | 0 | 10.7 | 7.52(4.12,10.91),1 | 9.92,1 | 9.49,1 |
| 34 | 2 | 86 | 1 | 0.2 | 4.73(0.68,8.78),0 | 6.25,0 | 5.97,0 |
| 35 | 2 | 64 | 1 | 1.8 | 6.95(2.37,11.54),0 | 9.18,0 | 8.78,0 |
| 36 | 2 | 63 | 1 | 2 | 7.07(2.4,11.75),0 | 9.34,0 | 8.93,0 |
| 37 | 2 | 71 | 0 | 2.2 | 6.15(1.96,10.34),0 | 8.12,0 | 7.77,0 |
| 38 | 2 | 67 | 0 | 2.6 | 6.6(2.22,10.97),0 | 8.71,0 | 8.33,0 |
| 39 | 2 | 51 | 0 | 3.3 | 8.72(2.28,15.17),0 | 11.52,0 | 11.02,0 |
| 40 | 2 | 70 | 1 | 3.6 | 6.26(2.03,10.49),1 | 8.26,0 | 7.9,0 |
| 41 | 2 | 72 | 0 | 3.6 | 6.05(1.89,10.2),1 | 7.98,0 | 7.63,0 |
| 42 | 2 | 81 | 1 | 4 | 5.17(1.13,9.21),1 | 6.82,1 | 6.52,1 |
| 43 | 2 | 47 | 0 | 4.3 | 9.36(1.98,16.73),0 | 12.36,0 | 11.82,0 |
| 44 | 2 | 64 | 0 | 4.3 | 6.95(2.37,11.54),1 | 9.18,0 | 8.78,0 |
| 45 | 2 | 66 | 0 | 5 | 6.71(2.28,11.15),1 | 8.86,1 | 8.48,1 |

Table 1 continued Prediction results and accuracy (last digit in predicted time: 0 inaccurate, 1, accurate)
| ID | stage | age | death | time | Median(95% CI) | MTTF | MPET |
| --- | --- | --- | --- | --- | --- | --- | --- |
| 46 | 2 | 74 | 1 | 6.2 | 5.84(1.73,9.94),1 | 7.7,1 | 7.37,1 |
| 47 | 2 | 62 | 1 | 7 | 7.2(2.43,11.97),1 | 9.51,1 | 9.09,1 |
| 48 | 2 | 50 | 0 | 7.5 | 8.88(2.22,15.54),1 | 11.73,1 | 11.22,1 |
| 49 | 2 | 53 | 0 | 7.6 | 8.42(2.38,14.47),1 | 11.13,1 | 10.64,1 |
| 50 | 2 | 61 | 0 | 9.3 | 7.33(2.46,12.2),1 | 9.67,1 | 9.25,1 |
| 51 | 3 | 49 | 1 | 0.3 | 5.83(2.41,9.24),0 | 7.69,0 | 7.36,0 |
| 52 | 3 | 71 | 1 | 0.3 | 3.97(2.19,5.75),0 | 5.24,0 | 5.01,0 |
| 53 | 3 | 57 | 1 | 0.5 | 5.07(2.68,7.45),0 | 6.69,0 | 6.4,0 |
| 54 | 3 | 79 | 1 | 0.7 | 3.45(1.56,5.34),0 | 4.55,0 | 4.35,0 |
| 55 | 3 | 82 | 1 | 0.8 | 3.27(1.32,5.23),0 | 4.32,0 | 4.13,0 |
| 56 | 3 | 49 | 1 | 1 | 5.83(2.41,9.24),0 | 7.69,0 | 7.36,0 |
| 57 | 3 | 60 | 1 | 1.3 | 4.81(2.68,6.94),0 | 6.35,0 | 6.07,0 |
| 58 | 3 | 64 | 1 | 1.6 | 4.48(2.58,6.39),0 | 5.92,0 | 5.66,0 |
| 59 | 3 | 74 | 1 | 1.8 | 3.76(1.96,5.56),0 | 4.97,0 | 4.75,0 |
| 60 | 3 | 72 | 1 | 1.9 | 3.9(2.12,5.68),0 | 5.14,0 | 4.92,0 |
| 61 | 3 | 53 | 1 | 1.9 | 5.43(2.6,8.27),0 | 7.17,0 | 6.86,0 |
| 62 | 3 | 54 | 1 | 3.2 | 5.34(2.63,8.05),1 | 7.05,0 | 6.74,0 |
| 63 | 3 | 81 | 1 | 3.5 | 3.33(1.4,5.27),1 | 4.39,1 | 4.2,1 |
| 64 | 3 | 52 | 0 | 3.7 | 5.53(2.56,8.49),1 | 7.3,1 | 6.98,1 |
| 65 | 3 | 66 | 0 | 4.5 | 4.33(2.49,6.17),1 | 5.71,1 | 5.47,1 |
| 66 | 3 | 54 | 0 | 4.8 | 5.34(2.63,8.05),1 | 7.05,1 | 6.74,1 |
| 67 | 3 | 63 | 0 | 4.8 | 4.56(2.61,6.51),1 | 6.02,1 | 5.76,1 |
| 68 | 3 | 59 | 1 | 5 | 4.89(2.69,7.1),1 | 6.46,1 | 6.18,1 |
| 69 | 3 | 49 | 0 | 5 | 5.83(2.41,9.24),1 | 7.69,1 | 7.36,1 |
| 70 | 3 | 69 | 0 | 5.1 | 4.11(2.32,5.89),1 | 5.42,1 | 5.19,1 |
| 71 | 3 | 70 | 1 | 6.3 | 4.04(2.26,5.82),1 | 5.33,1 | 5.1,1 |
| 72 | 3 | 65 | 1 | 6.4 | 4.41(2.54,6.27),1 | 5.82,1 | 5.56,1 |
| 73 | 3 | 65 | 0 | 6.5 | 4.41(2.54,6.27),1 | 5.82,1 | 5.56,1 |
| 74 | 3 | 68 | 1 | 7.8 | 4.18(2.38,5.98),1 | 5.52,1 | 5.28,1 |
| 75 | 3 | 78 | 0 | 8 | 3.51(1.64,5.38),0 | 4.63,1 | 4.43,1 |
| 76 | 3 | 69 | 0 | 9.3 | 4.11(2.32,5.89),0 | 5.42,1 | 5.19,1 |
| 77 | 3 | 51 | 0 | 10.1 | 5.63(2.52,8.73),1 | 7.43,1 | 7.11,1 |
| 78 | 4 | 65 | 1 | 0.1 | 1.69(0.77,2.61),0 | 2.23,0 | 2.14,0 |
| 79 | 4 | 71 | 1 | 0.3 | 1.52(0.7,2.34),0 | 2.01,0 | 1.92,0 |
| 80 | 4 | 76 | 1 | 0.4 | 1.4(0.61,2.18),0 | 1.84,0 | 1.76,0 |
| 81 | 4 | 65 | 1 | 0.8 | 1.69(0.77,2.61),0 | 2.23,0 | 2.14,0 |
| 82 | 4 | 78 | 1 | 0.8 | 1.35(0.56,2.13),1 | 1.78,0 | 1.7,0 |
| 83 | 4 | 41 | 1 | 1 | 2.57(0.3,4.84),0 | 3.4,0 | 3.25,0 |
| 84 | 4 | 68 | 1 | 1.5 | 1.6(0.74,2.47),1 | 2.12,1 | 2.03,1 |
| 85 | 4 | 69 | 1 | 2 | 1.58(0.73,2.42),1 | 2.08,1 | 1.99,1 |
| 86 | 4 | 62 | 1 | 2.3 | 1.78(0.78,2.79),1 | 2.35,1 | 2.25,1 |
| 87 | 4 | 74 | 0 | 2.9 | 1.44(0.65,2.24),0 | 1.91,1 | 1.82,1 |
| 88 | 4 | 71 | 1 | 3.6 | 1.52(0.7,2.34),0 | 2.01,1 | 1.92,1 |
| 89 | 4 | 84 | 1 | 3.8 | 1.21(0.42,2.01),0 | 1.6,0 | 1.53,0 |
| 90 | 4 | 48 | 0 | 4.3 | 2.28(0.57,3.99),1 | 3.01,1 | 2.87,1 |
| Accuracy rate(%) |  |  |  |  | 55.6%(50/90) | 50%(45/90) | 51.1%(46/90) |

## Discussion

In this paper, we introduced how to use a Weibull AFT model to predict when a health-related event will happen. Expected survival time, median survival time and minimum prediction error survival time from baseline to event were estimated and prediction accuracy assessed using Parkes’ and Christakis and Lamont’s method. When we fixed *k* = 2 the accuracy was 55.6% for median, 50% for MTTF and 51.1% for MPET. When used the method as suggested by Christakis and Lamont, the accuracy rate decreased to 30.0%,37.8 % and 33.3% (data not shown), respectively. Our example is limited as the sample is small and we only used two predictors. However, with a larger sample size and more predictors, accuracy may improve. In this example, we did not observe minimum prediction error time to be more accurate than median survival time.

Parametric survival models are advantageous in predicting survival time compared to semi-parametric Cox regression models. The Cox regression model which can be specified as 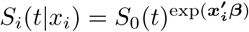 cannot predict time directly but calculates the probability of an event occurring within a specified timeframe. However, one disadvantage of parametric survival models such as Weibull AFT is the need to to make stronger assumptions than semi-parametric models. [14]

Currently, most clinical prediction models describe a patient’s likelihood of having or developing a certain disease as a traditional probability value or a risk score that is based on calculated probability. [15] However, probabilities are not intuitive to the general population and probability itself can be defined in different ways. [16] In practice, the time axis remains the most natural measure for both clinicians and patients. It is much easier to understand a survival time rather than probability of survival in a certain timepoint. Predicting when an event will happen provides a practical and interpretable guide for clinicians, health care providers and patients and and can help with decision making over remaining lifespan.

Upon recognizing that the Weibull AFT can be adapted from an engineering reliability framework to a medical framework, the next step involves developing a real prediction tool for medical predictive purposes, applying models to larger datasets and performing rigorous internal and external validation procedures. Such steps are outlined in the book Clinical prediction models: a practical approach to development, validation, and updating for the steps to develop a real prediction tool. [17]

